# BioBrigit, A Hybrid Deep Learning and Knowledge-based Approach to Model Metal Pathways in Proteins: Application to a Di-Copper Tyrosinase

**DOI:** 10.1101/2024.09.19.613875

**Authors:** Raúl Fernández-Díaz, Lorena Roldán-Martín, Mariona Sodupe, José-Emilio Sánchez-Aparicio, Jean-Didier Maréchal

## Abstract

The interaction of metallic species with proteins has been fundamental in evolution and key in many physiological processes. How metals bind to proteins also holds promise in many fields, like the design of new biocatalysts or the fight against pathogens. Nonetheless, uncovering the mechanism under which proteins recruit metal ions is far from understood and is one of the challenges in bioinorganic chemistry and structural biology. Computational methods are potentially among the most promising tools for this endeavor. Only a handful of efficient structural predictors of metal binding sites exist to date. Most focus on identifying the most stable binding sites in the protein scaffolds. Although these methods are very interesting, they do not consider the exploration of transient, sub-optimal binding sites that could be relevant in metal binding pathways in proteins. At the far end of modeling capabilities nowadays, we introduce BioBrigit, a hybrid Deep Learning – knowledge-based approach that suggests metal binding pathways in proteins. To demonstrate the method’s viability, we apply it to the di-copper tyrosinase from *Streptomyces castaneoglobisporus*, a system for which crystallographic experiments allowed the identification of a series of transient sites of the copper in its path from a chaperone to the final catalytic site. Combined with homology modeling and large-scale molecular dynamics, BioBrigit allows for computational characterization of all experimental sites and for better understanding of the copper recruitment mechanism. BioBrigit appears as an asset in a field full of unknowns like metal binding to proteins and opens the way to further algorithms in this area.

Source code, documentation, and data are available at https://github.com/insilichem/BioBrigit

## 1 Introduction

Incorporating metal ions in Nature’s molecular panoply has enabled a broad range of functions beyond the capabilities of organic frameworks.^1^ Transition metal ions that are parts of the biological landscape, like iron, copper, manganese, or zinc, are crucial for living organisms as they participate in processes like respiration, metabolism, or the stability of DNA motifs; but are also linked to diseases including neurodegenerative ones.^2^ Because of their relevance, computational approaches have soon been applied to decode the nature of the interplay between biological systems and metals.^3^

Several aspects of molecular modeling are still challenging in this field, particularly the prediction of metal binding sites in proteins and peptides. This information is crucial in many fields of modern chemistry, including elucidating the correct binding site in experimental or theoretical models of metalloproteins and the design of *de novo* metalloenzymes.^4^ Increasing efforts have been dedicated to developing metal binding site predictors in recent years. These include structure-based, sequence-based, and hybrid approaches that use genetic algorithms,^5^ statistical training^6^, or empirical models^6^ and machine learning (ML).^7^ While the prediction of the native metal binding site is becoming more feasible, some events, like the prediction of the metal’s recruitment pathway upon reaching the protein, are still underexplored.

Our first attempt at metal-binding site prediction came in the form of BioMetAll,^6,^ a software based on the pre-organization of the protein backbone, based on massive statistical analysis of existing structures of metalloproteins. Apart from the excellent yields of BioMetAll in identifying metal binding sites in X-ray structures, it also led to a series of byproducts. In our initial study, we observed that BioMetAll identifies non-optimal metal binding sites, which we interpreted as metal channeling sites in the route of the metal from the solvent to the catalytic site. Because studying metal recruitment by proteins is an orphan field, we decided to tackle this problem. Unfortunately, despite the accuracy of the first version of BioMetAll in finding native and transient metal binding sites, several physicochemical descriptors that exceed the first coordination sphere of the metal are not present in this algorithm. To solve this problem, we bet on a hybrid approach based on deep learning and statistical values while recovering a few concepts from BioMetAll.

In a way, considering the framework of deep learning (DL) and its contribution to molecular modeling, metal-binding site prediction could be considered a subset of ligand-binding prediction. Using convolutional neural networks over a 3D point cloud representation of the protein biochemical environment that explicitly considers different physicochemical properties has been proven quite successful in finding possible ligand-binding regions.^8^ The application of this algorithm to metal-binding sites has been explored, though, for a limited selection of metals and with an implementation that focuses on predicting what metal would be more likely to bind to a given protein region.^9^

Here, we present BioBrigit, a hybrid DL-statistics-based model for predicting the diffusion paths of metal in proteins. Added to the pure DL elements of the code, a metal-specific scoring function is set to improve the software’s versatility (e.g., it is valid for 28 different metals). We report the approach’s general idea, algorithmic core, benchmark for the prediction of metal binding sites (which appear comparable to current state-of-the-art ML-based software), and finally, how to apply it in predicting diffusion paths.

## 2 Material and Methods

In this section, we first report the BioBrigit algorithm and then provide details on other methods used for the showcase studies.

### 2.1 BioBrigit

#### 2.1.1 Algorithm Overview

The algorithm starts by creating a 1Å grid throughout the protein. Each point in the grid acts as a ‘probe’ that interrogates the surrounding protein environment to determine how likely it would be for a metal to occupy that position. This interrogation is a two-step process that relies first on an ML model that focuses more on finding general protein environments suitable for hosting metallic moieties and then evaluates those environments using a knowledge-based scoring function that provides more precise information regarding whether a metal ion could be coordinated by the protein at that location.

The algorithm is packaged within a command-line application and is implemented in Python.^10^ Its main dependencies are PyTorch^11^ and PyTorch Lightning for handling the ML component, and NumPy for handling the rest of the tensor operations.^12^ The rest of the dependencies are more specific and are discussed throughout the methodology when relevant.

#### 2.1.2 Machine Learning: CNN model

##### Architecture

The ML part of the algorithm consists of a convolutional neural network trained on 3-D point clouds (3D-CNN) representing protein environments to distinguish between metal-binding and non-binding regions ^8,9^. The model architecture is inspired by DeepSite^8^ and refined to the task at hand. Briefly, it contains a set of four layers with 32 neurons with (3 x 3 x 3) kernels, followed by a 3D Average Pooling layer, another set of four layers with 32 neurons with (3 x 3 x 3) kernels, and another 3D Average Pooling layer. Every two convolutional layers, a 3D Batch Normalisation layer is included. Finally, two linear layers with 512 and 256 neurons were included. All layers use leaky ReLU as a non-linear activation function. The final layer contains a single neuron with sigmoid function as its activation to provide an output in the range [0, 1].

##### Protein region representation

Protein environments are represented as point clouds with dimensions (12, 12, 12, 6) where the fourth dimension contains different physico-chemical properties that are associated with the traditional docking atom types, namely, i) hydrophobicity, ii) acceptance of H-bonds, iii) donation of H-bonds, iv) positive ionizability, v) negative ionizability, and vi) excluded volume. The representation of these properties as a point cloud is known as ‘voxelization’, and the program relies on the implementation of Moleculekit (Jiménez et al., 2017) to compute them.

##### Data set preparation

Protein-metal binding regions for 28 different metals (Li, Na, K, Rb, Cs, Be, Mg, Ca, Sr, Ba, V, Cr, Mo, Mn, Fe, Ru, Co, Rh, Ir, Ni, Pd, Pt, Cu, Ag, Au, Zn, Cd, and Hg) were extracted from the MetalPDB database.^13^ The benchmark used in Sánchez-Aparicio et al.^6^ containing 53 proteins with binding sites with the His-His-Asp/Glu motif was used as a hold-out evaluation set. Therefore, the 53 proteins were removed from the main training set. The rest of the structures were then filtered to remove those samples where the metal was not directly bound to the protein but to other entities (DNA, RNA, or other ligands). Further, those structures that either were not obtained through crystallographic methods or presented binding sites improperly annotated were also discarded. Negative samples were obtained by randomly selecting protein crystallographic structures from the PDB. A filter was implemented to consider only substructures with at least one amino acid present. This process was performed until the number of non-binding sites reached the number of metal-binding sites, so the training set was balanced. Ultimately, the training data set consisted of 103,530 sites from 30,186 proteins.

This data-gathering strategy implicitly under-sampled non-binding regions to avoid imbalance problems.^14^ Further, the combination of such a diverse array of metals within the binding data set was expected to direct the model towards recognizing second coordination features within the regions that allowed the identification of binding sites rather than first coordination elements that are much more heterogeneous across the periodic table.

##### Model training

The model was trained for 100 epochs with binary cross-entropy as loss function, ADAM optimizer, and early stopping for mitigating any possible overfitting. The training hyper-parameters were selected after prior experiments regarding optimal batch size (64) and learning rate (10-4). A strategy of 10-fold cross-evaluation was employed for training and robustly validating the models. More information about the evaluation metrics considered is included in the supplementary material.

#### 2.1.3 Knowledge-based scoring function

##### The concept

The scoring function estimates the probability that a certain probe will bind to the protein and is expressed as: *p*(*probe*). This probability is approximated by considering three different components computed for each residue-probe interaction:

1. **Metal-residue binding affinity:** certain metals are more likely to bind to specific residues. We model this affinity as the proportion of coordination bonds that the metal establishes with a residue in the MetalPDB database *N*(*metal ∩ residue*) over the total number of coordination bonds documented for that metal *N*(*metal*),

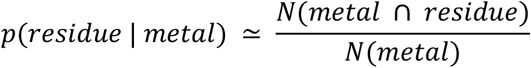
2. **Protein backbone pre-organization:** previous work from our research group (Sánchez-Aparicio et al., 2020) has showcased the importance of protein backbone pre-organization for forming protein-metal binding sites. Here, we expand upon this concept by creating a fitness function that measures how close a certain protein region’s pre-organization aligns with the ideal binding geometry. To approximate this ideal state, we performed a statistical analysis of three geometrical descriptors of coordinating residues, i.e., i) the distance between the metal and *C*_*α*_ (*α*), ii) the distance between the metal and *C*_*β*_ (*β*), and iii) the angle formed between both distances (*γ*). Our analysis was performed for each individual metal. We then approximated the probability density function of the distribution of each parameter as a bimodal function, both for its mathematical simplicity and for the shape of the distributions observed. Thus, the likelihood that a given residue can coordinate a certain metal, based on the set of geometrical parameters *x* = {*α, β, γ*}, is approximated as the union of the three distributions.

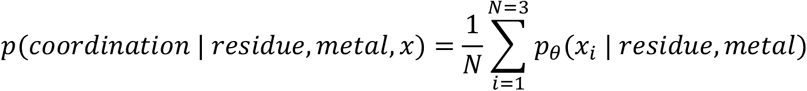

where each 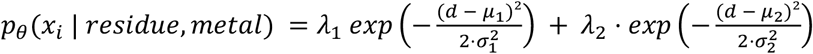 and is thus, defined by a set of 6 parameters *θ* = {*λ*_1_, *λ*_2_, *μ*_1_, *μ*_2_, *σ*_1_, *σ*_2_}. *μ*_*i*_ are the centres of the distributions; *σ*_*i*_ the deviations; and *λ*_*i*_ a correction factor that accounts for the relative contributions of each distribution component. To find the most appropriate set of parameters *θ*, the empirical data distributions were smoothened with a Kernel Density Estimator (KDE) with gaussian kernel^15^ and then the curves *p*_θ_ (*x*_*i*_ | *residue, metal*) were adjusted to the smooth distribution using the Levenberg-Marquardt algorithm for non-linear least-squares adjustment.
3. **Boolean filter:** To make the search for binding sites more efficient, a filter was implemented to discard unlikely regions easily. If one of the values for the geometrical descriptors falls out of 3 times the interquartile range of its distribution for that metal and residue, then the score is 0; else, it is 1.

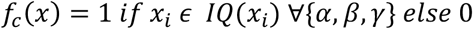 Combining the three expressions, we obtain the final scoring function:

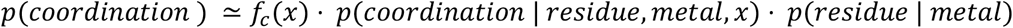

##### Data gathering

The data used for all the statistical analysis was obtained following the methodology described in the BioMetAll’s publication. However, the data was updated with the new structures that became available over the last year.

##### Integration of ML and Knowledge-based approaches and software availability

The main algorithm is illustrated in Figure 1. The first step is the voxelization of the protein. The mesh of points generated in this step will also be used as probes. A sliding window is used to evaluate the chemical environment of each probe with the CNN model. Those points with a CNN score superior to a certain threshold (by default: 0.5) are proposed for coordination analysis. The coordination analysis is the knowledge-based part of the approach and takes advantage of the algorithm described in [BioMetAll v2.0 inline citation]. The idea is to evaluate the availability of residues for directly coordinating the probe and assign a score to that possible coordination. The final score for each probe is calculated as the weighted sum between the CNN score and the coordination analysis score. This hybrid scoring function depends on a customizable weight (by default: 0.5), such that:

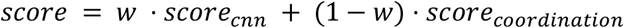

**Figure 1.**
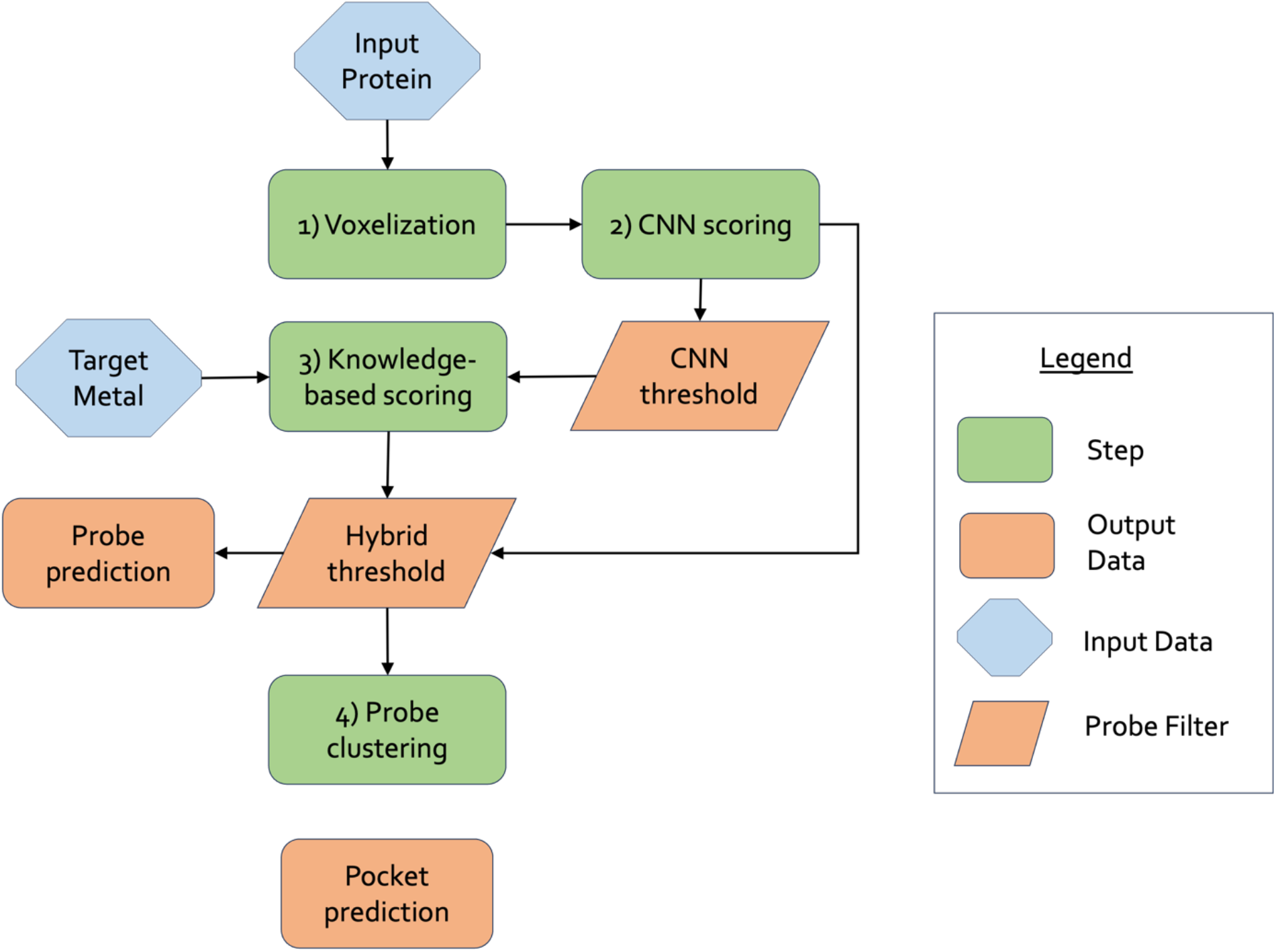
BioBrigit algorithm workflow.

BioBrigit, a command-line software was built using the Python programming language in order to use this model easily. All code and documentation are available at the Github repository: [https://github.com/insilichem/BioBrigit].

### 2.2 Computational details of other methods used in the study

#### 2.2.1 Initial Models for Tyrosinase

The caddie-tyrosinase (CTyr) complex crystal structure (PDB 3ax0),^16^ soaked with Cu(II) at different times, has been selected for the setting up of the computational approach. As the caddie subunit is incomplete in the crystallographic model, the missing loops were modelled with Homology Modelling in UCSF Chimera Software.^17^

#### 2.2.2 Molecular Dynamics simulations

CTyr structure was submitted to Classical Molecular Dynamics without Cu ions in the catalytic centre. The complexes were embedded in a cubic box of pre-equilibrated TIP3P water molecules and Na+ ions to balance the total charge according to the needs of each system. Simulations were performed with the AMBER ff14SB^18^ force field in the NPT ensemble for 300ns, using an integration time step of 1 fs. Constant temperature and pressure were set by coupling the system to a Monte Carlo barostat at 1.01325 bar and a Langevin thermostat at 300 K. The SHAKE algorithm was used to constrain hydrogen atom bonds.^19^

GaMD simulations^20^ were also performed in parallel, with coordinates starting after 10ns of MD simulations. The AMBER ff14SB force field in the NVT ensemble was applied, with SHAKE applied to constrain the bonds involving H atoms and an integration time step of 2 fs. A dual boost was performed on dihedral and total potential energy (igamd=3), finally obtaining a GaMD of 1μs long.

#### 2.2.3 Docking

Docking simulations were performed for the Tyrosinase system (PDB code 3ax0)^16^ in the areas predicted by the ML code to assess their suitability for metal binding. Protein-ligand dockings calculations were performed with the GOLD program at each high punctuation cluster,^21^ using the GoldScore scoring function, including the GOLD improvement to account for metal ions interaction and their possible coordinating amino acids.^22^ Genetic algorithm (GA) parameters were set to 50 GA runs and a minimum of 100,000 operations each. The remaining parameters were set to default, including pressure, number of islands, crossovers, and mutations. Finally, docking solutions were analysed through GaudiView^23^ an in-house developed GUI tool built as an extension of UCSF Chimera.^17^

#### 2.2.4 Analysis

To evaluate the exploration exhaustiveness, the MD simulations were subjected to Principal Component Analysis (PCA),24 RMSD all-to-all, and clustering.

## 3 Results

### 3.1 BioBrigit Capabilities - identification of primary and secondary metal binding sites

The ability of BioBrigit to find known primary metal-binding sites was evaluated by comparing against the BioMetAll hold-out benchmark set comprising 53 crystallographic structures of proteins bound to different transition metals (Zn, Fe, Mn, Ru, Co, Ni, and Hg). We considered a binding site as identified if any of the probes is situated below 3Å from the known crystallographic position of the metal ion. 69% of the binding sites were correctly located within the top-5 predicted binding sites with an average distance of 2.0 ± 1.0 A and 45% within the top-1 predictions. These results are notably inferior to the performance achieved by the original BioMetAll (98% and 75%, respectively); though still comparable to other state-of-the-art tools like IonCom^25^ and MIB.^7, 26^

A close examination of the predictions made by BioBrigit in target systems showed a greater ability to identify regions that, though not necessarily primary binding sites, presented interesting biochemical properties that could reasonably contribute to a transitory stabilization of metallic ions. The lack of proper literature describing metal migration phenomena makes it impossible to properly quantify the capability of BioBrigit to distinguish these putative processes. The best-case scenario for validating the predictiveness of BioBrigit would be to compare the computational outcomes with a series of metalloproteins with experimental structures (e.g., X-ray or NMR) displaying different snapshots of the metal intake process. Unfortunately, the protein data bank offers very little in this sense. In fact, to the best of our knowledge, the only system we could use as a benchmark is the copper tyrosinase from *S. castaneoglobisporus*.^4^ Therefore, we challenged BioBrigit to “re-”discover the metal recruitment path of this tyrosinase as observed in the X-ray structure and through the analysis of this particular use case, we introduce a qualitative evaluation of its ability for discovering transitory metal-binding sites.

The predictions from BioBrigit can be easily visualized with UCSF Chimera (or any other similar software) by using the B-factor of the probes generated to color them. This visualization is particularly relevant as it highlights one of the strengths of the combination of knowledge-based scoring and the CNN model. They can reconstruct continuous paths of metal-stabilizing regions from outside the protein to its final location. Through this path, there are regions predicted with lower affinity and certain points with high affinity. In the case of the tyrosinase model system case, as we will show presently, the points with high affinity correspond to secondary metal-binding sites confirmed through experimental and computational studies. The high-affinity regions are mostly predicted by the knowledge-based scoring functions, which are fitted to predict the first coordination sphere of the metallic moieties, and the lower-affinity regions are mostly predicted by the CNN. This suggests that the CNN has learned to identify elements that could stabilize the second coordination sphere, but it is not as precise for the first coordination sphere. If we look at the representation used (cubical regions of 12 Å in size), the volume they cover provides a coarse-grained approximation of the physicochemical properties surrounding the point of interest that would be compatible with the second coordination sphere. Regardless, a closer examination of this phenomenon is impossible without access to more data regarding metal migration paths.

### 3.2 Metal Diffusion in di-copper tyrosinases

Tyrosinases and tyrosinase-related enzymes are a very broad family of metalloenzymes whose members involve different kinds of cofactors: heme, di-copper, and di-zinc.^27^ Copper tyrosinases catalyze the hydroxylation of phenol and oxidation of catechol to quinone, finally producing dopaquinone, a melanin precursor (Scheme 1). Melanin is a widespread pigment in many species, from bacteria, fungi, plants, and mammals. Di-copper tyrosinases are found in almost all organisms, and mutations have been classically related to albinism and melanoma in humans.^28^ In contrast, neuromelanin, produced by the brain form of tyrosinase, has been recently linked to Parkinson’s Disease (PD).^29^ Strategies aiming at decreasing neuromelanin levels have been proposed as suitable therapeutic approaches for PD. Hence, tyrosinase inhibition and activation could be considered important targets.

**Scheme 1.**
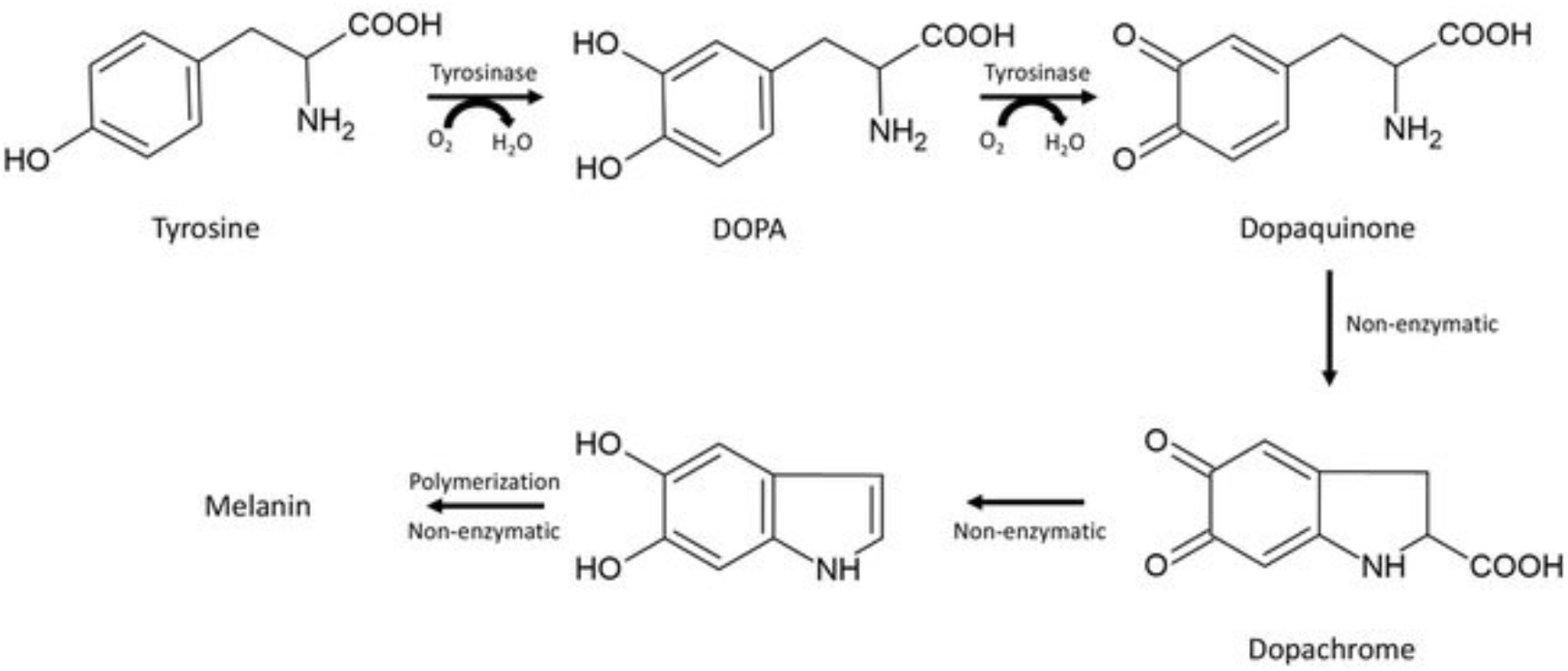
Melanin production from tyrosine. Tyrosinases are involved in the first two steps of the reaction, DOPA and dopaquinone production.

Copper tyrosinases have been shown to recruit copper in different ways. The *Streptomyces castaneoglobisporus* tyrosinase, however, cannot capture copper ions by itself and requires the aid of a metallochaperone, allowing the metal to transfer from the environment to the active site. In the tyrosinase system from *S. castaneoglobisporus*, the metallochaperone is called caddie, whose mutations have been observed to cause a lower activation of the tyrosinase. The tyrosinase-caddie complex is observable in the crystal published by Matoba et al.,^16^ with several Cu binding sites identified. Even though it is already demonstrated in the crystal that the caddie protein is actively involved in the process of Cu caption and transport to the inside of the tyrosinase, the initial transition site proposed for Cu(II) is already buried inside the chaperone, not exposed to solvent, suggesting that a first step is not observed in this structure. Nonetheless, the caddie crystallographic structure misses an external loop with high flexibility, solvent exposure, and remarkable histidine content, which could be crucial for Cu(II) recruitment. In this work, we rely on a computational pipeline based on BioBrigit to analyze the metal binding channel in *Streptomyces castaneoglobisporus* and compare experiments with calculated pathways.

In the work of Matoba et al.,^16^ five discrete binding locations for Cu(II) were identified in the CTyr crystal structure by soaking the protein at different times, revealing a well-defined metal recruiting channel (Figure 2). Two are in the active site and correspond to catalytic metal ions: i) Cu(A), being coordinated by His190T, His194T, and His216T, and ii) Cu(B), being coordinated by His38T, His54T, and His63T (T stands for Tyrosinase). Matoba *et al*. also describe three additional noncatalytic sites, which they name A, B, and C, and represent intermediate locations in the transition of the metal from the solvent-accessible region to the binding site. Site A is the most solvent accessible. There, Cu(II) is bound to residues Glu67C, His68C, and His82C, all belonging to the caddie protein (C stands for the Caddie protein). Site B is still entirely located inside the caddie, which requires the rotation of His82C, which is involved in Site A, to allow the coordination of Cu by both His82C and His97C. Site C represents the transition between the caddie and the tyrosinase, suggesting that the coordination requires the transport of the Cu ion by a rotameric movement of His97C, which would finally coordinate with His97C, His54T, and a water molecule. Interestingly, His54T is part of both the penultimate transition step (Site C) and the coordination sphere of the catalytic site for Cu(B).

**Figure 2.**
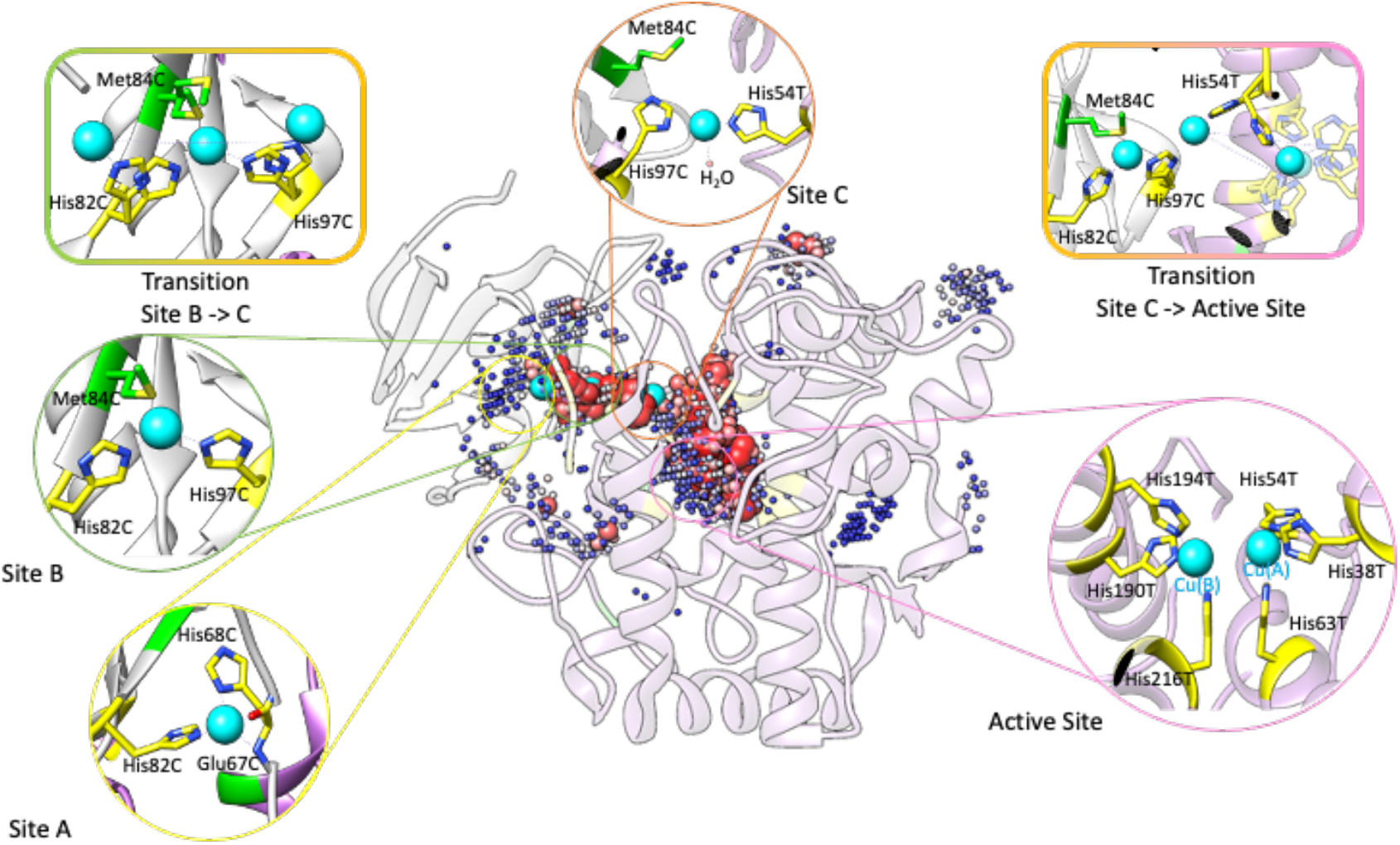
Binding sites observed in the crystallographic structure by Matoba et al.^16^(PDB code 3Ax0) overlapped with BioBrigit prediction for metal binding. Red spheres correspond to more likely binding sites (score of 1), while the blue ones correspond to less likely binding sites (score of 0.5).

A first calculation was carried out on the X-ray structure published by Matoba and coworkers.^16^ As expected, BioBrigit calculations correctly predict all catalytic and transient copper binding sites observed in the experimental structure (Figure 2). Interestingly, the predicted sites show differences in score between the more solvent-exposed regions, which present a score of 0.5 for each probe (blue in Figure 2), in the most exposed regions, such as Site C; while the more buried ones, such as the active site with the two catalytic coppers, scores of 1 (red in Figure 2) (see Table 1). In fact, the 5 best-punctuated regions by BioBrigit effectively correlate with the Active Site (Cu(A) and Cu(B)) (cluster score of 143.5 and 96.7), and detect the residues that comprise sites A (73.6 cluster score), B and C (122.5 and 101.9 cluster score, detecting transition areas). Other sites (clusters 6 to 14 in Table 1) with lower affinities (cluster score below 45) are scattered at the protein surface, which cannot be associated with any diffusion path, whose role is yet unknown, although alovalency could not be discarded.^16^

**Table 1.** BioBrigit cluster results found in S. *castaneoglobisporus*, with particular coordinators and scores of each cluster.

| Cluster | Coordinators | Score of the cluster |
| --- | --- | --- |
| 0 | His38T, His63T, His190T, His194T, His216T<br>(Active Site) | 143.5 |
| 1 | Met43T, Asp45T, His68T, His82C, Met84C<br>(Transition Site C -> Active Site) | 122.51 |
| 2 | His54T, Glu182T, His190T, Asn191T, His97C<br>(Transition Site C -> Active Site) | 101.88 |
| 3 | His38T, His54T, His63T (Active Site) | 96.7 |
| 4 | His66C (Site A) | 73.85 |
| 5 | Glu67C, His68C, Gln80C, His82C (Site A) | 59.6 |
| 6 | Glu175T, His180T, Arg185T | 44.34 |
| 7 | Arg229T, His230T, Asp232T | 37.07 |
| 8 | Asn6T, Asp87T, Asp91T | 22.9 |
| 9 | Arg228T, Asp264T, His265T, Tyr269T | 12.19 |
| 10 | Asp87T, Arg153T, Tyr218T, Lys221T | 10.18 |
| 11 | Arg140T, Glu114C | 10.12 |
| 12 | Arg185T, Asp88C | 7.59 |
| 13 | Glu149T | 7.53 |
| 14 | Asp112T, Tyr138T | 6.68 |

Though BioBrigit reaches excellent agreements with the metal binding sites observed in the X-ray structure, one of the remaining questions concerns identifying the first site for metal recruitment. Indeed, the first copper experimentally and theoretically observed is already buried inside the chaperone, suggesting that a first step may be missing. So, the question is whether another metal binding site is better exposed to the solvent.

Interestingly, some BioBrigit probes are detected in a slightly more solvent-exposed caddie area than in the X-ray structure, with low affinity, including residues from site A (cluster number 5). This hints at a possible additional site for metal recruitment as this region corresponds to part of the caddie’s metallochaperone missing in the X-ray structure and comprises two additional histidine residues (60 to 65 with sequence GGGAHH). Considering its flexibility, solvent exposure, and histidine content, such a missing region could be important for Cu(II) recruitment.

A Homology Model was built to complete the metallochaperone missing loop to address the first question further and study the caddie metal recruitment capacity at the solvent-exposed region using Modeller V5.0 plugin in UCSF Chimera.^30^ On this model, BioBrigit indeed predicts a metal binding site at the cluster of amino acids His298C, His299C, His300C, His302C, and His316C (Figure 3), which suggests that the X-ray misses a crucial part of the system in terms of metal binding. This new site (Table 2, cluster 0) allows us to understand the chaperone’s role better. It suggests that the first step in metal binding is a solvent-exposed, highly flexible, and histidine-rich region composed entirely of metallochaperone amino acids. Notably, the BioBrigit score at this new site is by far the highest (ca. 200 units versus ca. 120 for the ions of the catalytic sites)

**Table 2.** BioBrigit cluster results found in S. *castaneoglobisporus*, with specific coordinators and scores of each cluster.

| Cluster | Coordinators | Score of the cluster |
| --- | --- | --- |
| 0 | His298C, His299C, His300C | 217.2 |
| 1 | His298C, Glu301C, His302C, Gln314C, His316C | 84.6 |
| 2 | His316C, Met318C, His331C | 68.31 |
| 3 | His302C | 58.87 |

**Figure 3.**
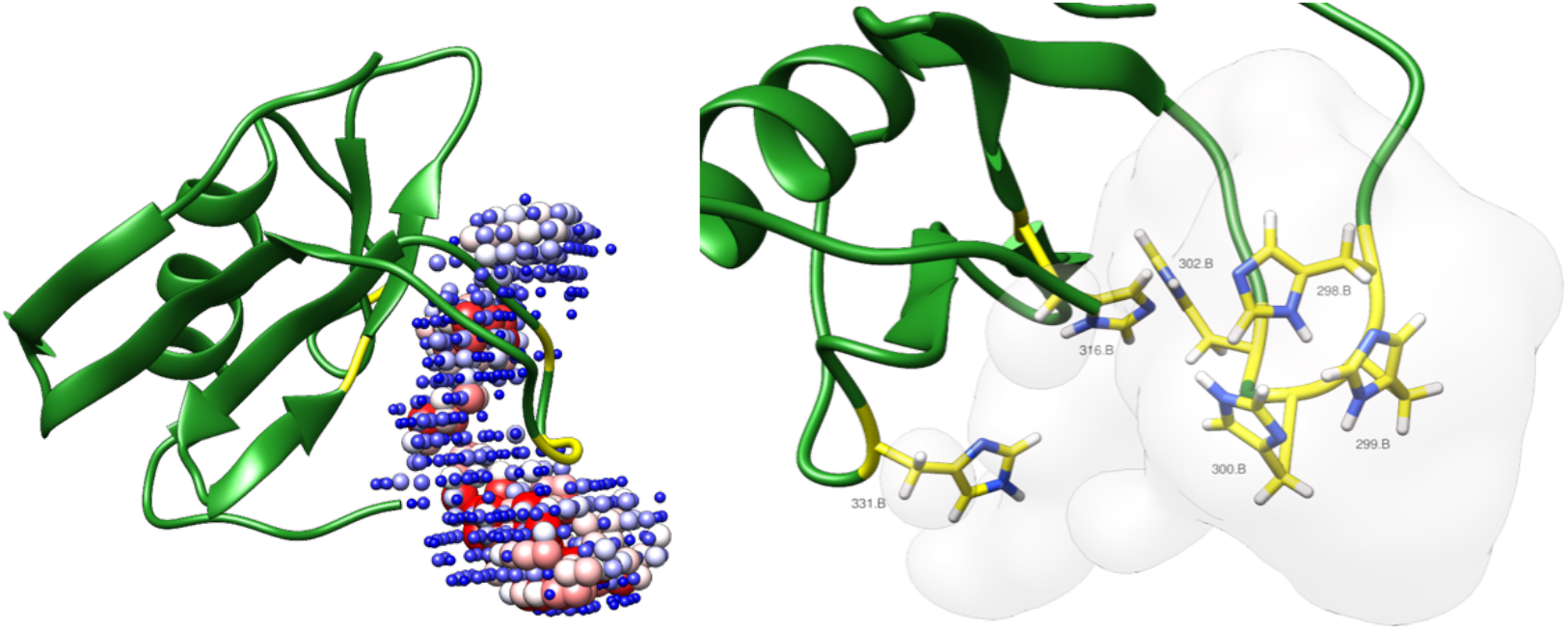
A. Metallochaperone model, with added missing regions from the X-ray structure. A cluster of His appears (yellow residues), which BioBrigit predicts as a high binder region.

However, BioBrigit only provides probes and does not describe the coordination geometry. To further demonstrate how it could be helpful in a more extensive exercise for modeling metal recruitment, metal-compatible protein-ligand dockings as set up in our group were performed for copper at sites 0 to 5 identified in the Tyrosinase and 0 in the metallochaperone, which show the higher scores (table 2).

Docking calculations reinforce the BioBrigit prediction with sites A, B, and C in the tyrosinase involving convenient orientation of the residues Glu67C, His68C, and His82C for A; His82C, Met84C, and His97C for B; and His97C, Asp45T and His54T for C (Figure 4). Besides, docking calculations reinforce site A’ in the chaperone as a feasible first step for metal recruitment. A critical aspect involves the necessity for particular residues, such as Glu67C, His82C, and His97C, to undergo reorientation to facilitate the transport of metal ions from one location to the next position. These dockings sustain our hypothesis about the rotameric transition of His97C. BioBrigit could not observe the transition between the chaperone and the tyrosinase since His97C is the only coordinated residue to Cu during the transition, which retrieves a meager score for the software.

**Figure 4.**
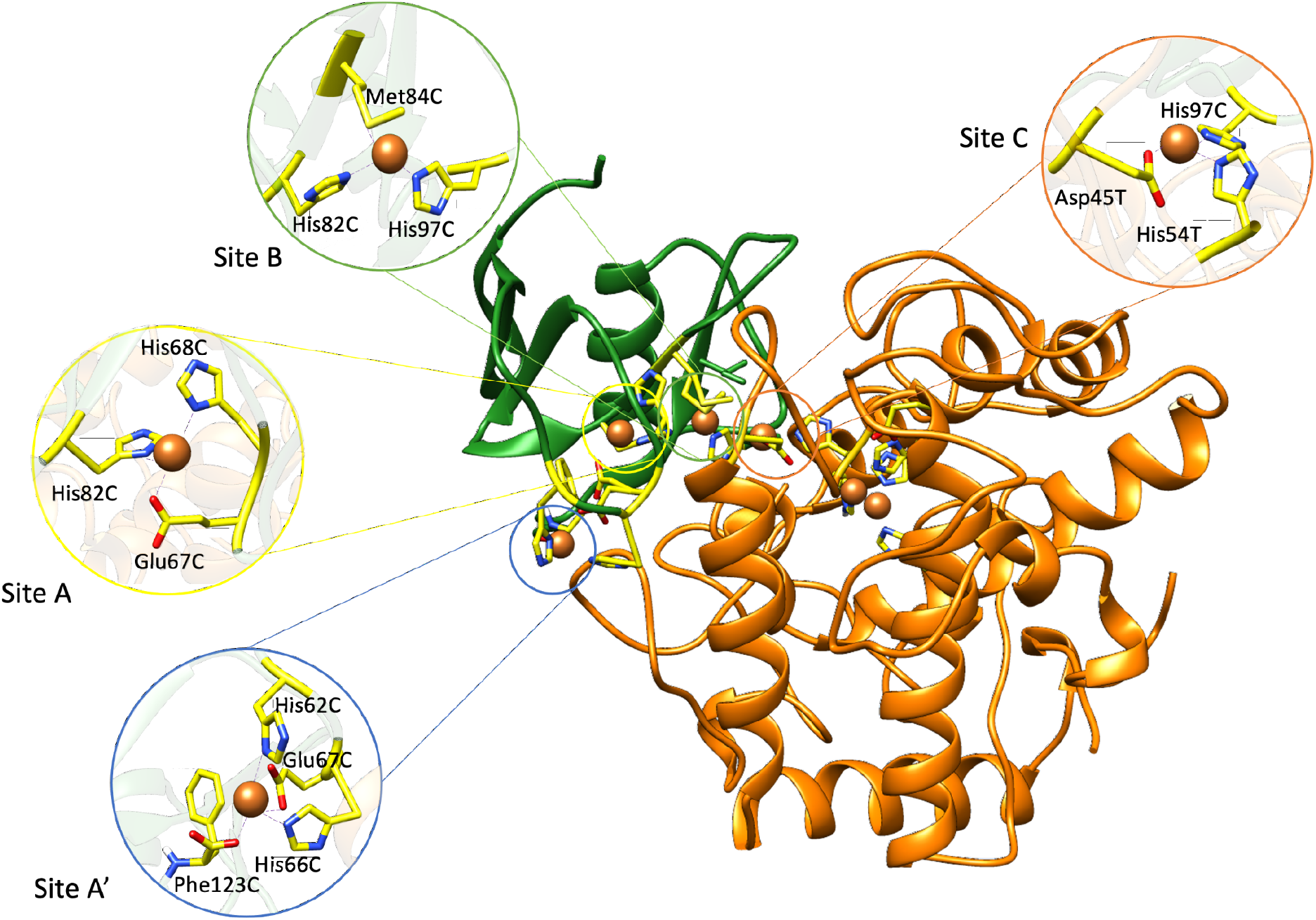
Docking results for sites A’, A, B, and C, which are very similar to the crystallographic determined ones (A, B, and C), and demonstrate that site A’ as feasible for metal binding. Glu67C, His82C and His97C are identified as links between each coordination site.

Although CTyr is known to be folded without metal, local rearrangements cannot be easily overruled. Based on the analysis performed on the X-ray structure, one could naturally wonder how copper recruitment is coupled with the receptor’s molecular motions. A GaMD simulation of the apo form of the CTyr complex was performed to assess this aspect.

The overall analysis of the 1μs GaMD simulations shows that the system is very stable, with no significant conformational change observed in the structure, except for the chaperone His-rich loop (His299C, His300C, His302C, His316C), which demonstrates high flexibility, as expected. A representative structure was obtained from the most populated cluster of the simulation to assess how dynamical events affect the copper recruitment path. BioBrigit calculations were carried out on this geometry. Results show some new, low-affinity areas (cluster score between 37 and 1) on the protein’s surface that do not lead to the active site. Hence, the apo structure and the crystal demonstrate the same metal pathway, starting in the His-rich loop of the metallochaperone, with no striking differences between the loaded states.

In summary, this part of the study on CTyr shows that i) BioBrigit effectively predicts the Cu-metal sites and channel as suggested by the X-ray structure, ii) several rearrangements of the amino acids from the X-ray structure are required to complete the path, for example Met84C, iii) the apo form of the complex is stable and maintain the same transition path for the metal, and iv) the missing region of the X-ray structure at position 60 to 65 is fundamental for the metal recruitment, which is absent in the experimental structure but it appears as the first step in metal recruitment.

## 4 Conclusion

Here, we present BioBrigit, a metal binding site predictor based on a hybrid ML-knowledge-based approach suitable for up to 28 different metals. Though producing results accurately in the range of its competitors regarding native metal binding, BioBrigit appears particularly relevant for transient sites and metal diffusion paths. The algorithm appears, therefore, to be a complementary tool in the interaction of metal ions with proteins.

The relevance of BioBrigit in this field is further demonstrated by creating a computational pipeline that includes Homology Modelling, Docking, Molecular Dynamics, and BioBrigit, and applying it to studying metal channels in bacterial tyrosinase. The channel, observed in the X-ray structure by Matoba et al.14, was successfully reproduced, showing the quality of BioBrigit for its prediction task. We also observed two additional results: 1) the detection of a high-affinity site absent in the crystal and likely initiator of the entire metal recruitment process and 2) the identification of geometric motions that should accompany the transition. The essential role of the caddie protein, a metallochaperone in charge of recruiting and transporting the metal ion to the entrance of the active site, has also been demonstrated. Because of the relevance of tyrosinase in the production of melanin and its relationship with melanoma in cancer, the correct identification of the metal recruitment process could be relevant for future biomedical and biotechnological applications.

Overall, both BioBrigit and the integrative pipeline outlined here have proven to be a valuable tool for systems reliant on metal ion loading for their functionality as they facilitate the effective identification of metal ion channels. Its applicability to other metalloproteins, particularly those with undiscovered druggable target sites, may represent a major advance in treating diseases.

## Acknowledgment

The authors thank the support of the Generalitat de Catalunya (2021 SGR-866) and the Spanish Ministerio de Ciencia e Innovación for Grant PID2020-116861GB-I00 and PID2023-149492NB-I00.

